# Virulence Differences of Monkeypox Virus Clades 1, 2a and 2b.1 in a Small Animal Model

**DOI:** 10.1101/2022.12.01.518711

**Authors:** Jeffrey L. Americo, Patricia L. Earl, Bernard Moss

## Abstract

Human monkeypox, a disease with similarities to smallpox, is endemic in Africa where it has persisted as a zoonosis with limited human-to-human spread. Unexpectedly, the disease expanded globally in 2022 driven by human-to-human transmission outside of Africa. It is not yet known whether the latter is due solely to behavioral and environmental factors or whether the monkeypox virus is adapting to a new host. Genome sequencing has revealed differences between the current outbreak strains, classified as clade 2b, and the prior clade 2a and clade 1 viruses but whether these differences contribute to virulence or transmission has not been determined. We demonstrate that the wild-derived inbred CAST/EiJ mouse provides an exceptional animal model for investigating clade differences in monkeypox virus virulence and show that the order is clade 1 > clade 2a > clade 2b.1. The greatly reduced replication of the clade 2b.1 major outbreak strain in mice and absence of lethality at 100-times the lethal dose of a closely related clade 2a virus, despite similar multiplication in cell culture, suggest that clade 2b is evolving diminished virulence or adapting to other species.

**SIGNIFICANCE:** Three clades of monkeypox virus are recognized: clade 1 is present in the Congo Basin, causes 10% human mortality and is transmitted by rodents with little human-to-human spread; clade 2a exists in West Africa, has a low mortality and is also a zoonosis; clade 2b is currently spreading globally by human transmission. The genetic basis for differences in virulence and transmission have not been determined. A major roadblock is the need for a small animal model that can be studied under the stringent safety conditions required. Here we demonstrate that the three clades exhibit highly significant differences in CAST/EiJ mice in the order clade 1 > clade 2a > clade 2b, similar to the severity of clinical disease.

## INTRODUCTION

Although first isolated from captive Asian monkeys, monkeypox virus (MPXV) infects rodents and is incidentally transmitted to humans and non-human primates in Africa (1). Human monkeypox, a disease resembling smallpox, was recognized upon the eradication of the latter and the cessation of vaccination (2). Historically, monkeypox has been a zoonosis with little human-to-human transmission and until 2022 was largely limited to regions of Africa. The genome sequence of the Congo Basin (now referred to as clade 1) MPXV (3) placed it in the orthopoxvirus genus of the chordopoxvirus subfamily along with variola virus, the causative agent of smallpox as well as vaccinia virus used as the smallpox vaccine. A second clade of MPXV (now referred to as clade 2a) with a 95% nucleotide sequence identity to the original was subsequently recognized (4). Clade 1 is associated with a mortality of up to 10% whereas clade 2a from West Africa has a mortality of less than 1%. The current outbreak strain of MPXV, has been designated clade 2b because of its sequence similarity to clade 2a (5) and like the latter has a low mortality in immunocompetent individuals. Whereas clades 1 and 2a are zoonoses, clade 2b has exhibited extensive human-to-human spread. Although aerosol transmission of MPXV has been demonstrated in the laboratory (6), direct contact with animals and close human interactions appear more important (7).

The number of human monkeypox cases has been increasing in Africa, likely due to waning population immunity from the discontinued smallpox vaccine as well as changing environmental and social factors (8). Human monkeypox due to Clade 2a has occurred in other parts of world through importation of African rodents into the U.S. in 2003 and by travelers from Africa but without evident human-to-human transmission (9). However, the unprecedented increase in clade 2b cases in 2022 is due to human transmission outside of Africa. At the time of writing, more than 80,000 cases in over 100 different locations were diagnosed.

Genetic differences contributing to the greater morbidity of clade 1 MPXV compared to clade 2a and the increased human transmission of clade 2b are unknown. Knowledge of such differences could provide an early warning of the potential virulence of new or hybrid strains, help identify therapeutic targets, and contribute to basic knowledge of virus-host interactions. Although the genomes of clade 1 and 2 viruses are highly conserved, there are numerous sequence differences that could account for the greater virulence of the former (4, 10), whereas clades 2a and 2b viruses have fewer differences (5, 11). Animal models are crucial for investigating virus pathogenesis, and in the case of MPXV such studies need to be carried out under stringent BSL-3 containment conditions. Moreover, clade 1 viruses are classified as Select Agents in the US limiting the number of laboratories working on MPXV. Parker and Buller (12) and Alakunle et al. (13) have provided extensive descriptions of natural and experimental infections with MPXV, although the primary reservoir species is unknown. The experimental animals include Asian primates, African rodents, and a variety of North American rodents including prairie dogs and squirrels. The African dormouse (Graphiurus kelleni) is of particular interest since these small squirrel-like rodents can be bred in captivity and were among the African rodents that tested positive for MPXV in a shipment that introduced the virus into the US in 2003. Cotton rats (Sigmoidin sp.) are another species susceptible to MPXV that can be raised in captivity. Nevertheless, African dormice and cotton rats are not inbred, and the absence of immunological reagents is a disadvantage for the study of virulence. Other small animals that can be reared in captivity such as guinea pigs, hamsters, rats, and most mouse strains do not develop severe disease upon inoculation with MPXV.

A mouse model for MPXV would have numerous advantages. To identify a susceptible mouse, we screened a panel of 38 inbred strains by intranasal (IN) inoculation of a Clade 1 MPXV and discovered the exceptional vulnerability of the CAST/EiJ mouse and confirmed the resistance of common inbred mouse strains (14). Subsequent studies showed that the CAST mouse is also highly susceptible to other orthopoxviruses including vaccinia, cowpox and Akhmeta viruses (15, 16). IN inoculation of CAST mice with variola virus, the causative agent of smallpox, resulted in only mild disease with some virus shedding and limited skin lesions (17) in accordance with the specificity of variola virus for humans.

CAST mice were derived from founder specimens trapped in a grain storage facility in Thailand (JAX^®^ NOTES, Issue 456, Winter 1994) and because they are genetically distinct from common inbred laboratory mice were included in the Collaborative Cross panel (18). CAST mice are immunologically competent; they make good antibody and T-cell responses upon vaccination that provide complete protection from MPXV (14). Low basal levels of natural killer cells likely contribute to their susceptibility to MPXV and other orthopoxviruses, as administration of interferon gamma, IL-15 or natural killer cells expanded *in vitro* each provide resistance (15, 19-21). Here, we develop the CAST mouse as a model to compare MPXV clades and show that the order of virulence is clade 1 > 2a > 2b.1.

## RESULTS

### Clade 1 MPXV and clade 2a MPXV lethally infect African dormice

Schultz and co-workers (22) reported that African dormice are highly susceptible to MPXV by the intranasal (IN) route with a median 50% lethal dose (LD_50_) of 12 PFU for the clade 1 MPXV-ZAI-1979-005, and they cited a similar though undisclosed value for MPXV Copenhagen-58, a 2a clade. We confirmed and extended these studies using clonal isolates of MPXV-ZAI-1979-005, abbreviated here as Z-1979, and clade 2a MPXV-USA-2003-044 (abbreviated USA-2003). No difference in replication of the two viruses was found during multi-step growth in African green monkey BS-C-1 cells (Fig. 1A). Upon IN infection of African dormice with Z-1979, some deaths occurred with 5 and 50 PFU and all died with 500 and 5,000 PFU (Fig. 1B). IN infection with USA-2003 followed a similar pattern with 100% deaths with 5,000 PFU (Fig. 1C). LD_50_ values of 12 and 5 PFU were calculated for Z-1979 and USA-2003, respectively. Large numbers of non-inbred African dormice would be required to establish biological significance of the relatively small difference in LD_50_ and therefore were not considered suitable for investigating clade differences.

**Fig. 1.**
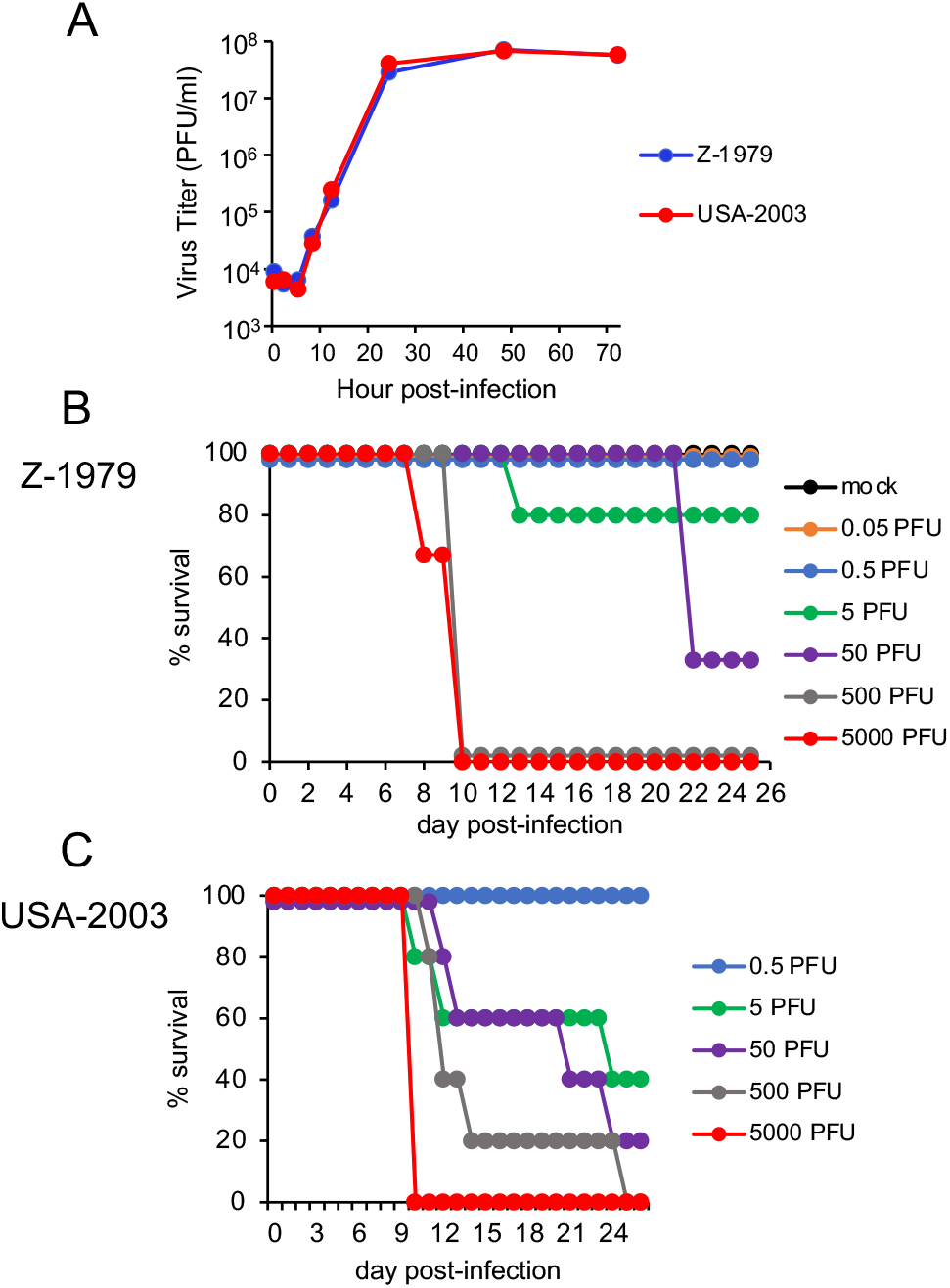
MPXV Z-1979 and USA-2003: *in vitro* replication and African dormouse infection. **(A)** BS-C-1 cells were infected in triplicate with 0.05 PFU/cell of MPXV Z-1979 or USA-2003. At intervals virus yields were determined by plaque assay on BS-C-1 cells. Bars indicate standard deviation. **(B)** Each group (n=3-5) of African dormice were mock infected or infected IN with 0.05 to 5,000 PFU of Z-1979 and survival plotted. **(C)** Same as panel B except animals were inoculated with 0.5 to 5000 PFU of USA-2003.

### Clade 1 MPXV is up to 1,000 times more virulent than clade 2a MPXV in CAST mice

CAST mice infected IN with 10^3^ PFU of Z-1979 lost more than 20% of their weight and one died (Fig. 2A). At 10^4^ and 10^5^ PFU all mice infected with Z-1979 died or were euthanized because of 30% weight loss. Following IN infection with USA-2003, comparable weight loss occurred at slightly higher doses and most mice died at 10^4^ PFU and all at 10^5^ PFU. From these data, LD_50_ of 2,370 and 4,200 were obtained for Z-1979 and USA-2003, respectively. In addition, there was greater recovery of virus from the nasal turbinates, lungs, brain, and abdominal organs on days 5 and 7 after IN infection of Z-1979 compared to USA-2003 (Fig. 2B).

**Fig. 2.**
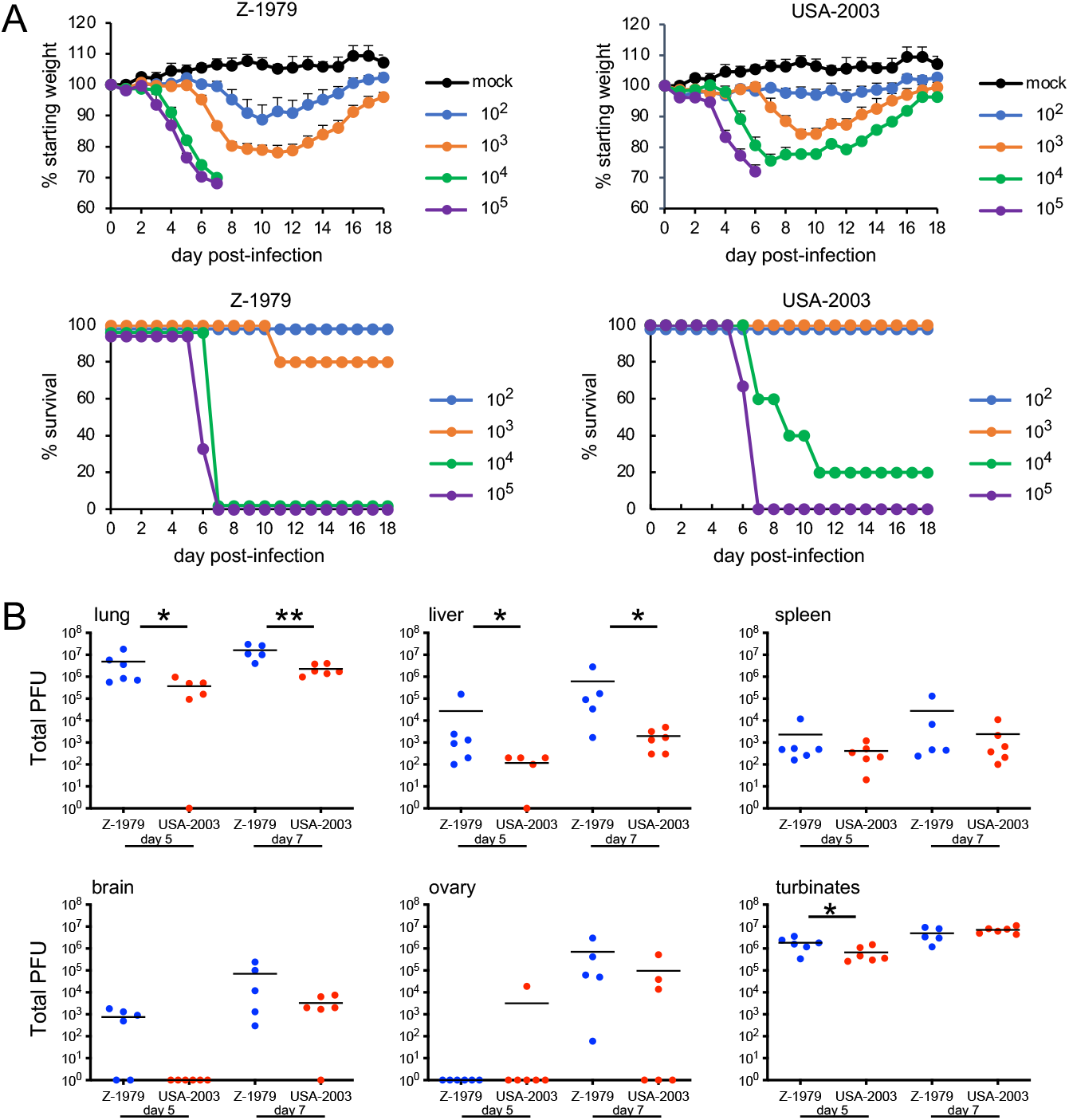
Virulence of MPXV Z-1979 and USA-2003 upon IN infection of CAST mice. **(A)** CAST mice (groups of n=4-5) were mock infected or infected IN with 10^2^ to 10^5^ PFU of MPXV Z-1979 or USA-2003 and examined and weighed daily. The percent of starting weights and survival are indicated. Bars indicate standard deviation. **(B)** CAST mice (n=4-5) were infected IN with 10^2^ PFU of MPXV Z-1979 or USA-2003. On days 5 (d5) and 7 (d7), organs were removed, homogenized and the total PFU determined by plaque assay. * p<0.05, ** p<0.01).

Since the upper respiratory tract is not thought to be an important route of MPXV infection and extensive spread of virus to abdominal organs occurred by day 5 after IN infection, we investigated a systemic route of infection. Remarkably, most CAST mice infected with Z-1979 died with 1 and 10 PFU and all with 100 and 1,000 PFU administered intraperitoneally (IP), despite little weight loss (Fig. 3A). In contrast, USA-2003 caused no deaths at 100 PFU and only 50% succumbed to 1,000 PFU. The LD_50_ values were <1 for Z-1979 and 1,000 for USA-2003. Furthermore, the titers of the Z-1979 were significantly higher in all organs analyzed at day 6 after infection compared to USA-2003 (Fig. 3B). Note that mortality is greater by IP infection, but weight loss is more pronounced by IN infection, possibly because the latter route reduces eating and drinking.

**Fig. 3.**
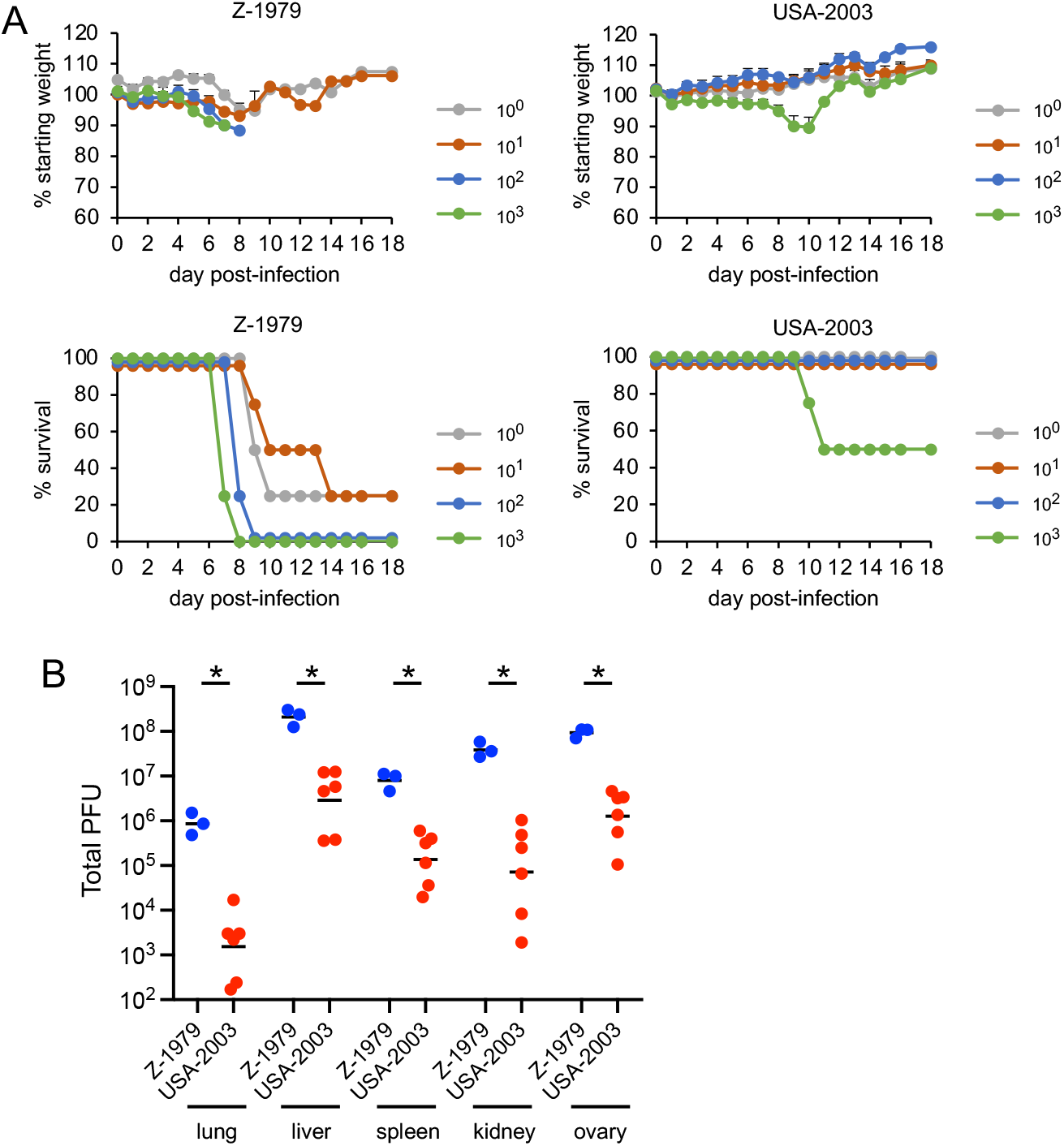
Virulence of MPXV Z-1979 and USA-2003 upon IP infection of CAST mice. **(A)** CAST mice (groups of n=4-5) were mock infected or infected IP with 1 to 10^3^ PFU of MPXV Z-1979 or USA-2003 and examined and weighed daily. The percent of starting weight and survival are indicated. Bars indicate standard deviation. **(B)** CAST mice (n=3-6) were infected IP with 10^2^ PFU of MPXV Z-1979 or USA-2003. On day 6, organs were removed, homogenized and total PFU determined by plaque assay. * p<0.05.

### MPXV clade 2b is more attenuated than clade 2a

The above experiments were carried out with MPXV isolated decades before the global 2022 outbreak due to MPXV clade 2b. We received a low passage MPXV isolated in 2022 from an individual in Massachusetts that is designated MPXV-USA-2022-MA001 (abbreviated MA-2022) and is a prototype for the major outbreak variant called b.1. Replication of the USA-2003 and MA-2022 at low (0.05 PFU/cell) and high (3 PFU/cell) multiplicities in African green monkey BS-C-1 cells was not significantly different, although there was a trend for greater release of USA-2003 into the medium compared to MA-2022 (Fig. 4A, B). Upon IN infection of CAST mice USA-2003 caused severe disease with weight loss and deaths at doses of 10^4^ and 10^5^ PFU (Fig. 4C), similar to that obtained for USA-2003 in the prior experiment shown in Fig. 2. In contrast, neither weight loss nor disease occurred with MA-2022. Furthermore, following IP infection USA-2003 caused weight loss and 100% lethality at 10^3^ and 10^4^ PFU, whereas neither weight loss nor lethality occurred at doses up to 10^5^ PFU of MA-2022 (Fig. 4D).

**Fig. 4.**
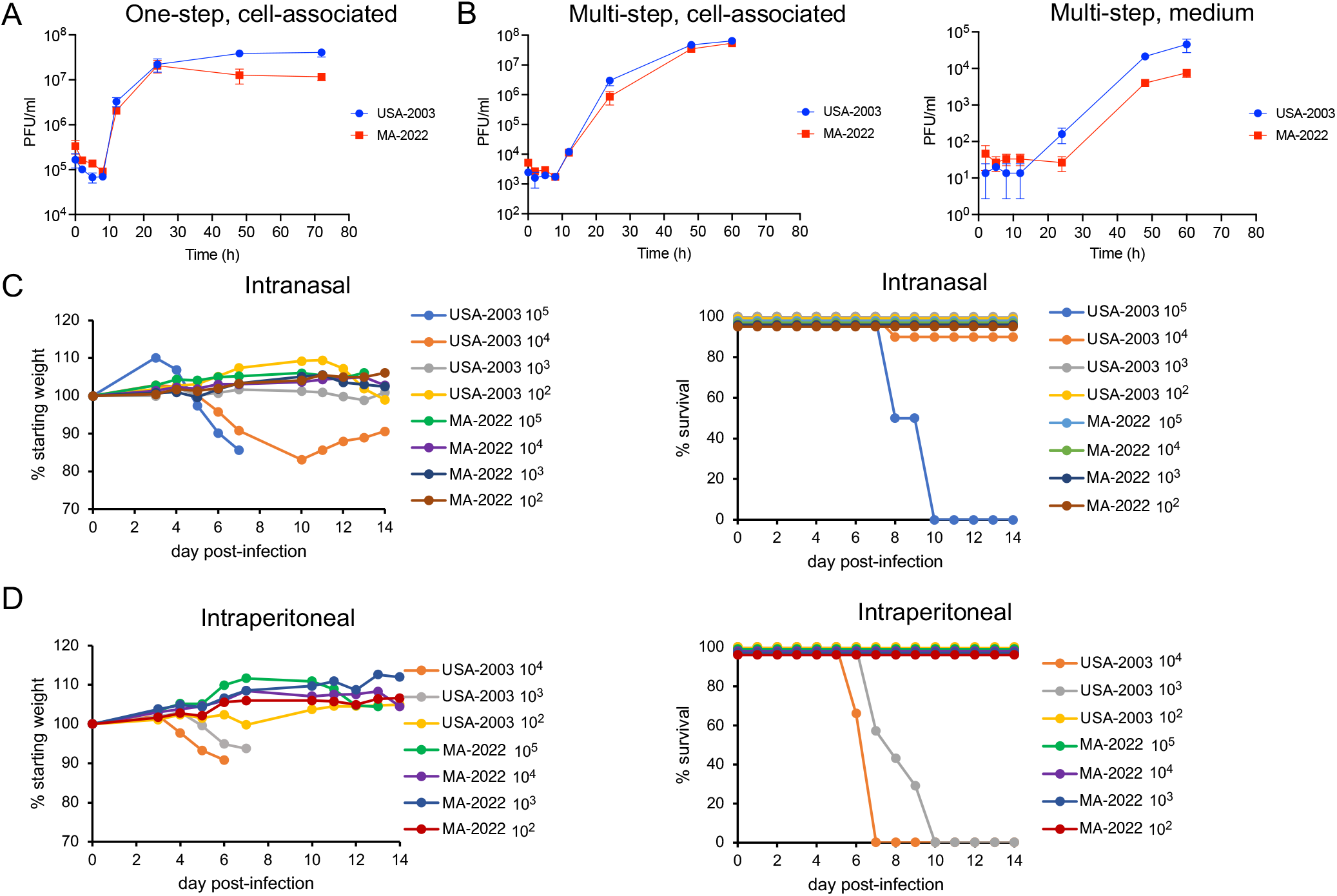
*In vitro* replication and virulence of USA-2003 and MA-2022. **(A)** One-step replication. BS-C-1 cells were infected in triplicate with 3 PFU/cell of USA-2003 or MA-2022 and harvested at the indicated times. Infectious virus in cell lysates was determined by plaque assay on BS-C-1 cells. Bars represent standard deviations. **(B)** Multi-step replication. BS-C-1 cells were infected in triplicate with 0.05 PFU of USA-2003 or MA-2022. At the indicated times, the medium was removed, centrifuged and supernatant collected. The pelleted cells were combined with the cells that adhered to the dish and lysed. Infectious virus in the lysates and supernatant were determined by plaque assay on BS-C-1 cells. Bars indicate standard deviations. **(C)** IN infection of CAST mice. CAST mice were inoculated with 10^2^ PFU (n=3), 10^3^ PFU (n=4) 10^4^ PFU (n=6-7) and 10^5^ PFU (n=2) of USA-2003 and MA-2022. Daily weights and survival are plotted. **(D)** IP infection of CAST mice. CAST mice were inoculated with 10^2^ PFU (n=3), 10^3^ PFU (n=4-7) and 10^4^ PFU (n=6) of USA-2003 and MA-2022. Daily weights and survival are plotted.

Next, we determined virus titers and genome copies in the organs of mice infected with 10^4^ PFU of USA-2003 and MA-2022. At 7 days after IN infection, the titers of USA-2003 were >10^6^ PFU and >10^5^ PFU in the lungs and nasal turbinates, respectively, and about 10^4^ PFU in the liver and spleen (Fig. 5A). In contrast, only 1 of 3 mice infected with MA-2022 had detectable virus in the lungs, none had detectable virus in liver or spleen and the mean titer in the nasal turbinates was 3 logs lower than that of USA-2003. Those results were consistent with an analysis of genomic DNA in the same samples by digital droplet (dd) PCR (Fig. 5B). We also determined virus titers and genomic DNA following IP infection with USA-2003 and MA-2022. The titers of USA-2003 were more than 100-fold higher than that of MA-2022 (Fig. 5C) and again this was corroborated by analysis of genomic DNA by ddPCR. Thus, the lower virulence of MA-2022 in CAST mice correlated with lower virus replication.

**Fig. 5.**
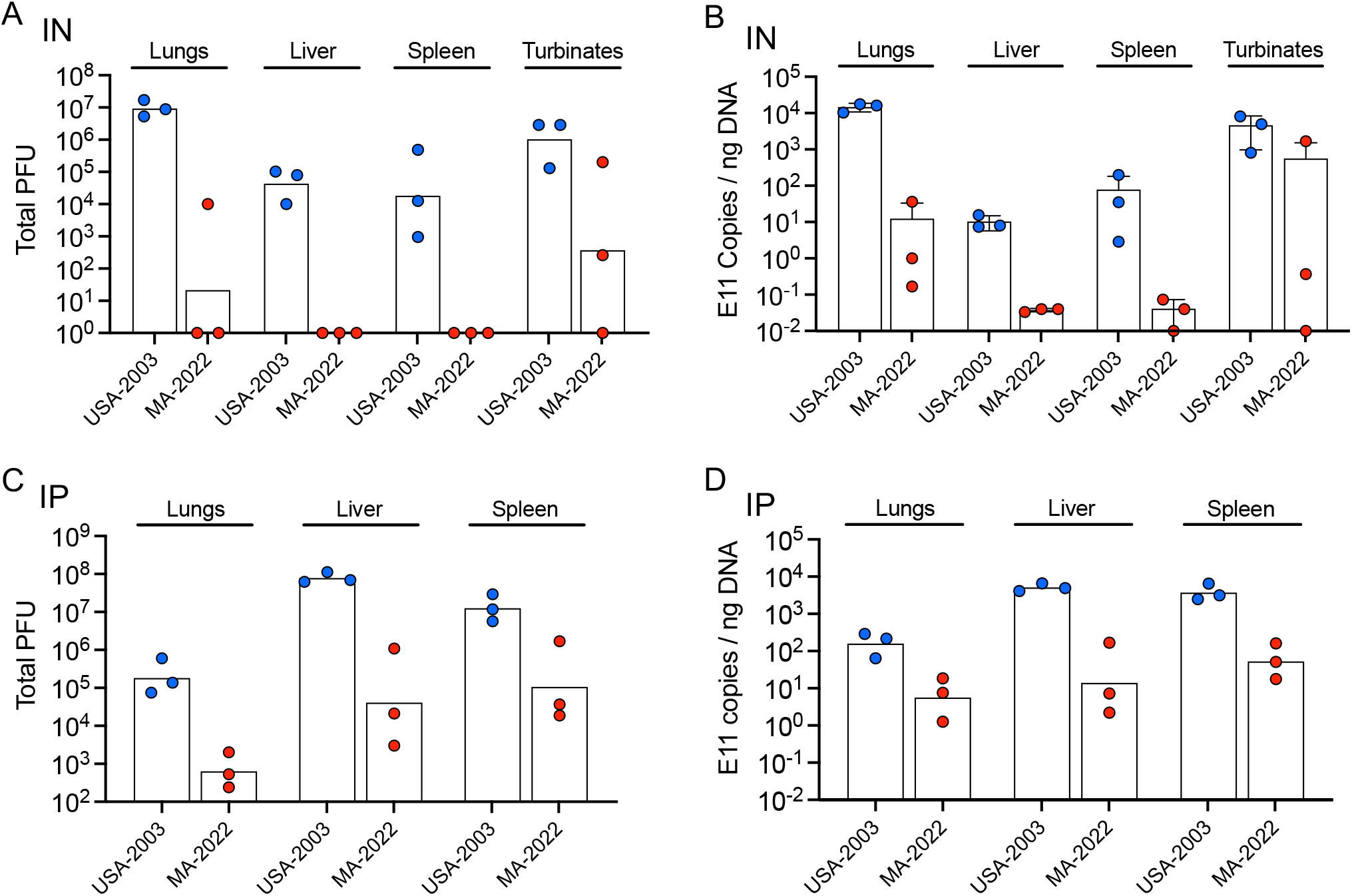
Infectious virus and genome copies of USA-2003 and MA-2022 in organs of CAST mice. **(A, B)** CAST mice (groups of n=3) were infected IN as in Fig. 5C and infectious virus and MPXV genome copies determined. **(C, D)** CAST mice (groups of n=3) were infected IP as in Fig. 5D and infectious virus and MPXV genome copies determined.

## DISCUSSION

There are numerous reports suggesting that observed genome sequence variations might account for differences in virulence and transmission of MPXV clades. However, few such predictions have been tested experimentally. Reasons for this include the need for a suitable animal model and the stringent biosafety conditions that must be met, particularly for clade 1 MPXV. Using objective criteria, we showed that clade 1, clade 2a and clade 2b.1 MPXVs exhibit large differences in morbidity and virus replication in CAST mice. By the IP route of infection, MPXV clades 1 and 2a had LD_50_ values of 1 PFU and 10^3^ PFU, respectively, whereas all mice survived even 10^5^ PFU of the clade 2b.1 virus. The difference in virulence between clades 1 and 2a was less by the IN route but here too mice survived 10^5^ PFU of the clade 2b.1 virus. For both routes, replication in the lungs and abdominal organs also followed the order of clade 1 > 2a > 2b.1. Although we have not yet compared clades using other routes for infection of CAST mice, we previously reported that skin scarification of the clade 1 virus causes local lesions but no other signs of disease (14), whereas footpad inoculation caused swelling of the lower leg and one of four mice died on day 14 with virus in the lungs, liver and spleen (14). Similar studies with clade 2a and 2b viruses are planned. The increase in human transmission of clade 2b and the striking reduction in virulence for CAST mice raise the possibility that MPXV is adapting to humans with a concomitant loss of fitness for certain other species.

Determination of which genes are responsible for the virulence differences of MPXV clades remains to be determined. MPXV like other members of the orthopoxvirus genus encode about 100 accessory genes that are largely dispensable for replication *in vitro* but affect host range and virulence. Interestingly, recent gene loss has been a major factor in the evolution of orthopoxviruses that can account for differences in their host range (23). For example, cowpox virus with the most accessory genes has a wide host range, whereas variola virus with the fewest is specific for humans. Of interest, human transmitted variola virus (17) and MPXV clade 2b.1 cause only mild disease in CAST mice. We have started to investigate the genetic determinants responsible for virulence differences of clade 1 and 2a viruses and plan to extend this to clade 2b. It has been suggested that orthopoxvirus immune modulators might reduce an overreaction of the host immune response and loss of such genes might increase virulence (24). Thus, both gain and loss of gene function should be considered in assessing virulence.

## METHODS

### Ethics statement

All procedures with infectious MPXV were performed in registered Select Agent BSL-3 and ABSL-3 laboratories by trained and smallpox vaccinated investigators using protocols approved by the NIH Institutional Biosafety Committee and judged not to have the potential for dual use research of concern (DURC). Experiments and procedures using mice were approved by the NIAID Animal Care and Use Committee according to standards set forth in the NIH guidelines, Animal Welfare Act and U.S. Federal law. Euthanasia was carried out using carbon dioxide inhalation in accordance with the American Veterinary Medical Association guidelines (2013 Report of the AVMA panel on euthanasia).

### Cells

African green monkey BS-C-1 cells (ATCC CCL-26) were maintained at 37°C and 5% CO_2_ in modified Eagle minimal essential medium (Quality Biological, Inc.) supplemented with 8% fetal bovine serum, 2 mM L-glutamine, 10 U penicillin/ml, and 10 µg streptomycin/ml.

### Viruses

Low passage stocks of MPXV-ZAI-1979-005, MPXV-USA-2003-044 and MPXV-USA-2022-MA1 were obtained from the CDC and procedures for isolation of clones of the 1979 and 2003 viruses have been reported (14). The MPXV Z-1979 clone expressing Luc (previously called MPXV-z06) has been described (25) and the same procedure was used to insert Luc into a clone of MPXV USA-2003. The primers used for construction of chimeric viruses are in the Extended Data Note.

### Analysis of virus replication

BS-C-1 cells in 12-well plates were infected with MPXV. After 1 h, monolayers were washed three times and overlayed with fresh medium. At various times, cells from triplicate wells were harvested and stored at -80°C. Cells were lysed by three rounds of freeze-thawing and samples were sonicated, and virus yields determined by plaque assay on BS-C-1 cells (26). For determination of extracellular virus, the medium was collected before harvesting cells, centrifuged and the supernatant analyzed.

### Mice

Female CAST/EiJ mice were obtained from Jackson Laboratories (Bar Harbor, ME) and were maintained in small, ventilated microisolator cages. IN and IP inoculations were performed as described previously (14). Titers of each virus inoculum were confirmed by plaque assay.

### Titration of virus from organs

Organs were removed aseptically from animals, placed in 2 ml of balanced salt solution supplemented with 0.1% bovine serum albumin, and stored at -80°C until used. Homogenization and titration were performed as described previously (14).

### Quantitation of viral DNA

Viral DNA from infected organs was determined by ddPCR as previously described (27). Briefly, DNA was isolated from homogenized MPXV-infected tissues using the DNeasy Blood and Tissue Kit (Qiagen). Several 10-fold dilutions of each purified DNA were prepared in nuclease-free H_2_O and added to a reaction mixture containing ddPCR EvaGreen Supermix (Biorad). Forward (5′-GAATACATTCACATTGACCAATCAGAA-3′) and reverse (5′GGTTCGTCAAAGACATAAAACTCATT-3′) primers were specific for the conserved VACV WR E11L open reading frame (27). Reaction oil droplets were synthesized using an Automated Droplet Generator (Biorad) prior to PCR thermocycling. Droplets were subsequently acquired using a QX200 Droplet Reader (Biorad) and the number of genome copies were determined using QuantaLife software (Biorad).

### Statistics

Significance was calculated by Mann-Whitney using Prism.

## ACKNOWLEDGEMENTS

We thank Catherine Cotter for assistance with some experiments and discussions. Excellent animal care was provided by the technical staff of the NIAID Comparative Medical Branch. The work was funded by the Division of Intramural Research, NIAID.

## REFERENCES

1. J. B. Doty et al., Assessing monkeypox virus prevalence in small mammals at the human-animal interface in the Democratic Republic of the Congo. Viruses-Basel 9 (2017).

2. K. Simpson et al., Human monkeypox - After 40 years, an unintended consequence of smallpox eradication. Vaccine 38, 5077–5081 (2020).

3. S. N. Shchelkunov et al., Analysis of the monkeypox virus genome. Virology 297, 172–194 (2002).

4. A. M. Likos et al., A tale of two clades: monkeypox viruses. J Gen Virol 86, 2661–2672 (2005).

5. C. M. Gigante et al., Multiple lineages of monkeypox virus detected in the United States, 2021-2022. Science 10.1126/science.add4153, eadd4153 (2022).

6. C. L. Hutson et al., Transmissibility of the monkeypox virus clades via respiratory transmission: Investigation using the prairie dog-monkeypox virus challenge system. Plos One 8 (2013).

7. A. T. Fleischauer et al., Evaluation of human-to-human transmission of monkeypox from infected patients to health care workers. Clinical Infectious Diseases 40, 689–694 (2005).

8. P. Y. Nguyen, W. S. Ajisegiri, V. Costantino, A. A. Chughtai, C. R. MacIntyre, Reemergence of human monkeypox and declining population immunity in the context of urbanization, Nigeria, 2017-2020. Emerging Infectious Diseases 27, 1007–1014 (2021).

9. M. R. Mauldin et al., Exportation of monkeypox virus from the African continent. Journal of Infectious Diseases 225, 1367–1376 (2022).

10. N. H. Chen et al., Virulence differences between monkeypox virus isolates from West Africa and the Congo basin. Virology 340, 46–63 (2005).

11. J. Isidro et al., Phylogenomic characterization and signs of microevolution in the 2022 multi-country outbreak of monkeypox virus. Nature Medicine 10.1038/s41591-022-01907-y.

12. S. Parker, R. M. Buller, A review of experimental and natural infections of animals with monkeypox virus between 1958 and 2012. Future Virology 8, 129–157 (2013).

13. E. Alakunle, U. Moens, G. Nchinda, M. I. Okeke, Monkeypox Virus in Nigeria: Infection Biology, Epidemiology, and Evolution. Viruses 12 (2020).

14. J. L. Americo, B. Moss, P. L. Earl, Identification of wild-derived inbred mouse strains highly susceptible to monkeypox virus infection for use as small animal models. J. Virol. 84, 8172–8180 (2010).

15. J. L. Americo et al., Susceptibility of the wild-derived inbred CAST/Ei mouse to infection by orthopoxviruses analyzed by live bioluminescence imaging. Virology 449, 120–132 (2014).

16. C. N. Morgan et al., Laboratory infection of novel Akhmeta virus in CAST/EiJ mice. Viruses-Basel 12 (2020).

17. N. F. Gallardo-Romero et al., Use of live Variola virus to determine whether CAST/EiJ mice are a suitable surrogate animal model for human smallpox. Virus Res 10.1016/j.virusres.2019.197772, 197772 (2019).

18. D. W. Threadgill, G. A. Churchill, Ten years of the collaborative cross. Genetics 190, 291–294 (2012).

19. P. L. Earl, J. L. Americo, B. Moss, Lethal monkeypox virus infection of CAST/EiJ mice is associated with a deficient interferon-gamma response. J. Virol. 86, 9105–9112. (2012).

20. P. L. Earl, J. L. Americo, B. Moss, Insufficient innate immunity contributes to the susceptibility of the castaneous mouse to orthopoxvirus infection. J. Virol. 91 (2017).

21. P. L. Earl, J. L. Americo, B. Moss, Natural killer cells expanded in vivo or ex vivo with IL-15 overcomes the inherent susceptibility of CAST mice to lethal infection with orthopoxviruses. PLoS Pathog 16, e1008505 (2020).

22. D. A. Schultz, J. E. Sagartz, D. L. Huso, R. M. L. Buller, Experimental infection of an African dormouse (Graphiurus kelleni) with monkeypox virus. Virology 383, 86–92 (2009).

23. T. G. Senkevich, N. Yutin, Y. I. Wolf, E. V. Koonin, B. Moss, Ancient gene capture and recent gene loss shape the evolution of orthopoxvirus-host interaction genes. mBio 10.1128/mBio.01495-21, e0149521 (2021).

24. A. Alcami, G. L. Smith, A mechanism for inhibition of fever by a virus. Proc. Natl. Acad. Sci. USA 93, 11029–11034 (1996).

25. P. L. Earl, J. L. Americo, C. A. Cotter, B. Moss, Comparative live bioluminescence imaging of monkeypox virus dissemination in a wild-derived inbred mouse (Mus musculus castaneus) and outbred African dormouse (Graphiurus kelleni). Virology 475, 150–158 (2015).

26. C. A. Cotter, P. L. Earl, L. S. Wyatt, B. Moss, Preparation of cell cultures and vaccinia virus stocks. Curr. Protoc. Mol. Biol. 117, 16 16 11–16 16 18 (2017).

27. J. L. Americo, P. L. Earl, B. Moss, Droplet digital PCR for rapid enumeration of viral genomes and particles from cells and animals infected with orthopoxviruses. Virology 511, 19–22 (2017).

